# Noninvasive focal transgene delivery with viral neuronal tracers in the marmoset monkey

**DOI:** 10.1101/2023.08.09.552625

**Authors:** T. Vincenza Parks, Diego Szczupak, Sang-Ho Choi, David J. Schaeffer

## Abstract

Although preclinical neuroscientific modeling species permit invasive intracranial delivery of targeted neurotropic agents, direct intracranial injections are not readily translatable to clinical therapeutics. Transcranial focused ultrasound (tFUS) has been identified as a technique to circumvent surgical injections altogether by transiently opening the blood-brain barrier (BBB) with selective focus. We have recently characterized the ability to focally deliver substances across the BBB in the marmoset, a non-human primate model with similar husbandry requirements to rodents but with cortical topologies more similar to humans. Here, we establish a reliable method for selectively delivering adeno-associated viral vectors (AAVs) across the BBB in marmoset frontal cortex with tFUS and demonstrate long-range anterograde neuronal tracing. Using a single-element 1.46 MHz transducer, we focally perturbed the BBB (∼1 x 2 mm) in area 8aD of frontal cortex in four adult marmoset monkeys using low-intensity focused ultrasound aided by microbubbles. We confirmed BBB opening via a gadolinium-enhanced MRI at 9.4 T prior to AAV delivery. Within an hour of opening the BBB, either AAV2 or AAV9 was delivered systemically via tail-vein injection. Four to six weeks later, animals were sacrificed, and microscopy was performed to confirm the presence of neurons transduced as indicated by EGFP or mCherry fluorescence. In all four marmosets, neurons were observed at the site of BBB perturbation, with AAV2 showing an exiguous distribution of transduced neurons when compared to AAV9. The results are compared to direct intracortical injections of anterograde tracers into area 8aD and similar (albeit sparser) long-range connectivity was observed. With evidence of transduced neurons specific to the region of BBB opening as well as long-distance tracing, we establish a framework for focal noninvasive transgene delivery to the marmoset brain. This technique will be of utility for the burgeoning marmoset model, with applications for noninvasive delivery of therapeutics, genetic delivery of precursors for techniques like two-photon imaging, or neuronal tracing across the lifespan.

## Introduction

Although preclinical neuroscientific modeling species allow for invasive intracranial delivery of neurotropic substances, direct intracortical injections are not readily translatable to clinical therapeutics. Recently, the use of transcranial focused ultrasound (tFUS) has been identified as a technique for circumventing direct injections, allowing for delivery of therapeutic across the blood-brain barrier (BBB) ^1^. Of particular interest for nonhuman primate species — who have a more similar genetic makeup to humans ^2^ – is the use of adeno-associated viruses (AAVs) as a vector for delivering neurotropic substances to the brain. As a small primate, the common marmoset is quickly becoming a premier neuroscientific modeling species ^3–5^. Marmosets have similar body size, housing and handling requirements as rodents, but with a brain with granular layers and connectivity more similar to humans ^6–10^ and a lissencephalic cortex (and very thin skull) that is more amenable to tFUS ^1^. Here, we establish the ability to noninvasively and focally deliver AAVs across the marmoset BBB with tFUS aided by microbubbles, allowing for anterograde axonal tracing.

Intracranial injection techniques have been fundamental for scientific progress in preclinical animal research, with much of our understanding of how the marmoset brain is structurally connected based on direct intracortical injections into marmoset brain parenchyma ^11–18^. Although direct intracortical surgical injections can offer exquisite targeting accuracy in well-trained hands, the requirement of trephination is accompanied by inherent risks of infection, long recovery times, and often concomitant tissue damage and behavioral complications. When considering therapeutic neurotropic injections, like gene therapies for human brain diseases, the requirement of transcranial surgery does not translate well to the clinic, especially when repeated administrations are required. Several approaches are being developed to circumvent the requirement for surgical injection by alerting the permeability of substances across the BBB. The use of the osmotic agent mannitol ^19^ and highly engineered AAV variants ^20,21^ are both promising methods for systemic agent delivery across the BBB. Although BBB crossing and neuronal transduction specificity can be achieved with AAV capsid variants ^20^, this noninvasive method currently only allows brain-wide transgene expression. Focal, mechanical perturbation of the BBB with tFUS aided by microbubbles allows for much more selective delivery – especially as required for techniques like neuronal tracing.

Recently we extensively characterized the use of tFUS to focally deliver substances across the BBB in marmosets ^1^. While optimizing for safety and limiting tissue damage, we established a reliable method for BBB opening with a single element 1.46 MHz transducer, resulting in detectable BBB extravasation volume of ∼1 mm radially and ∼2.5 mm axial. Indeed, this parenchymal volume is similar to that achieved through a direct nano-injection of viral tracer in marmosets ^18^. Although a relatively new technique, tFUS-mediated AAV delivery has been demonstrated in rodents ^22–24^ and more recently in macaque monkeys for the purpose of delivering viral vectors to subcortical regions involved in Parkinson’s disease ^25^. Here, we sought to build on these results to demonstrate that this technique is effective in New World primates (*Callithrix jacchus*) for the purpose of neuronal tracing, leveraging our recent advances in highly focal cortical BBB perturbation. Indeed, the reports in mice, rats, and macaques showed very sparse neuronal transduction and/or were delivered over a BBB opening across a large swath of cortex ^23,25,26^.

Here, we demonstrate long-range anterograde neuronal tracing from neurons transduced with either AAV2 or AAV9 with spatial specificity limited to the site of BBB perturbation, induced by tFUS. This technique required no opening of the cranial cavity and thus the focal delivery of the AAV to the brain was noninvasive. Four adult marmosets underwent low-intensity tFUS to focally open the BBB in area 8aD of frontal cortex. After confirming that the BBB was open using gadolinium-based contrast agent (GBCA) enhanced MRI at 9.4 T, AAV serotype 2 or 9 with the human synapsin promoter was injected into the tail vein. Four to six weeks later, ex vivo microscopy was performed to confirm the presence of neurons transduced as indicated by enhanced green fluorescent protein (EGFP) or mCherry fluorescence. We demonstrate long-distance connections, comparing the presence of axonal fluorescence to that of direct intracortical injections from the publicly available whole-brain marmoset tracer resource ^18^. With evidence of a sparse (AAV2) or more robust (AAV9) transduction of neurons in all four marmosets, as well as long-distance tracing, we establish a framework for which AAVs can be delivered to the marmoset brain noninvasively. This technique will be invaluable for the burgeoning marmoset model, with applications for noninvasive delivery of therapeutics, genetic delivery of precursors for techniques like two-photon imaging, or neuronal tracing across the lifespan.

## Results

### In vivo MRI-based assessment of BBB disruption

Following the sonication of right area 8aD 8/9/23 9:17:00 AM, each marmoset was immediately injected with a gadolinium-based contrast agent (GBCA) via tail catheter and transferred to the MRI. Using a magnetization prepared – rapid gradient echo (MPRAGE) or fast low angle shot (FLASH) sequence sensitized to the T1 relaxation times of gray matter with gadolinium ^1^ BBB disruption was confirmed after each sonication. The accuracy with reference to our target was confirmed with this same scan, as well as the extent of opening (Figure 1). Based on our previous work, the extent of GBCA extravasation as observed by MRI overlaps well with histological assessments of Evans blue extravasation ^1^. As such, the volume of BBB opening was also computed based on the GBCA contrast and used to localize our histological assessment of viral transduction. All but one animal required only a single sonication to open the BBB, with marmoset T requiring a second attempt with a slightly higher pressure and microbubble dosage (i.e., marmoset T was ‘resonicated’ because GBCA extravasation was not observed on the MRI). As shown in Figure 1, the relative size and location was consistent across animals – note that these sonications were atlas-based ^27^ and thus did not account for individual anatomical differences.

**Figure 1:**
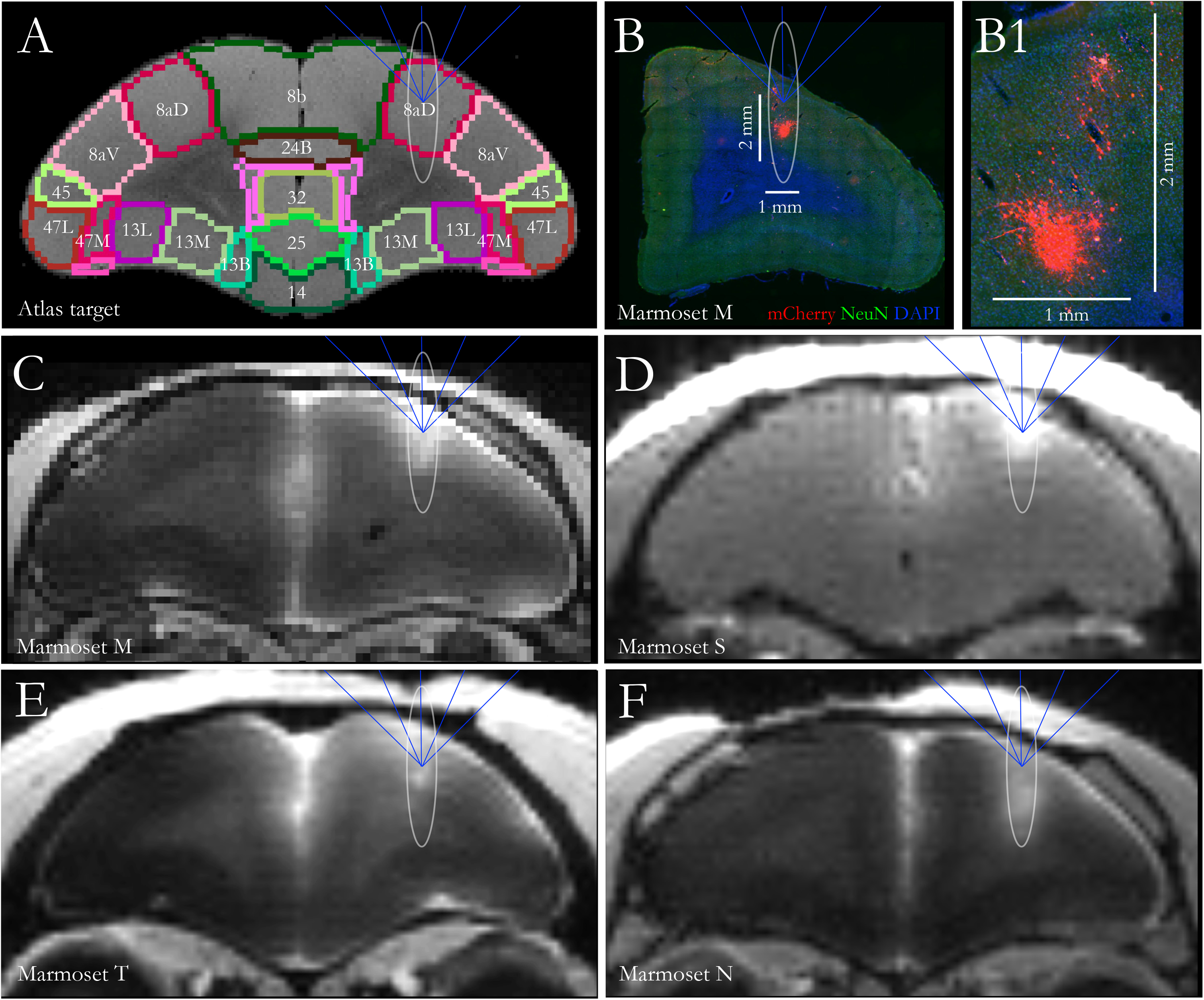
Atlas-based 8aD tFUS target, and resultant sonication sites for all four marmosets. A) Marmoset atlas-based target (8aD) which was integrated into the MORPHEUS (FUSInstruments) software. Ellipse shows the target volume of opening from the 1.46 MHz transducer – the full width at half maximum of the acoustic pressure distribution at the focus. B) Sonication target ellipse overlayed on slice from Marmoset M, showing both targeting accuracy and highly focal transduction as evidenced by mCherry florescence. B1) Zoomed view of the extent of viral transduction; labeled cells present in a 1 x 2 mm section of tissue. C-F) Gadolinium-enhanced MRI of Marmosets M (C), S (D), E (T), and N (F) showing successful blood-brain barrier perturbation prior to injection of AAV. Note Marmoset S had a FLASH image acquired rather than MPRAGE like the other 3 marmosets.

### Immunohistochemical assessment of viral transduction at the sonication site

Table 1 shows the serotype, titer, volume, and time to perfusion after BBB opening and intravenous delivery of AAVs. For both the animals receiving AAV9 (Marmoset S, AAV9-EGFP; Marmoset M, AAV9-mCherry) and those injected with AAV2 (Marmosets T & N, AAV2-EGFP), labelling was observed at the site of injection (Figure 1 for locations). Importantly, the observed expression from neuronal cell bodies was limited to parenchyma at the site of opening, as shown in Figure 2 (AAV9) and Figure 3 (AAV2), demonstrating the extent of focus possible with tFUS-AAV delivery with a 1.46 MHz transducer. See Supplementary Figure 1, however, for the atypical examples of non-specific transduction; note that Marmoset S (systemically injected with AAV9) showed the most observed nonspecific uptake of all the marmosets tested. When compared to AAV9 (Figure 2), the AAV2-mediated neuronal expression (Figure 3) was dramatically less. Although in many cases (especially with the sparse AAV2-mediate transduction) neurons could be unambiguously identified, triple immunolabelling for the EGFP or mCherry transgene with NeuN and DAPI allowed for identification of transduced neurons and nonneuronal cells, as shown in Figures 2 and 3. When comparing the four marmosets, neither serotype seemed to show laminar-level specificity, with transduced neurons observed across the depth of cortex.

**Figure 2:**
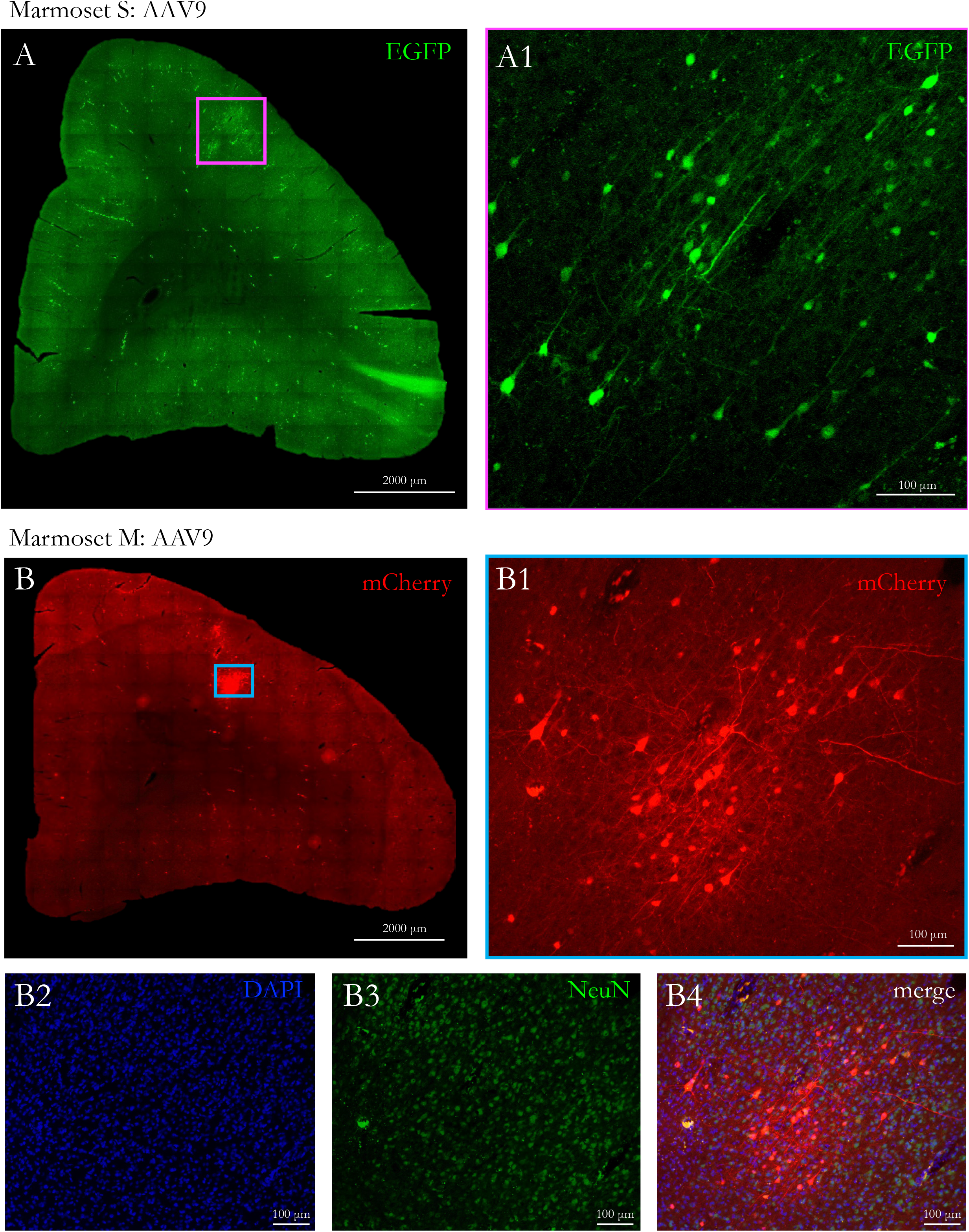
Neurons transduced at the sonication point in animals systemically injected with AAV9. A) EGFP-transduced neurons at sonication site in Marmoset S, right hemisphere 8aD. Scale bar: 2000 μm. Zoomed in view in A1 (scale bar: 100 μm). B) mCherry-transduced neurons at sonication site in Marmoset M, right hemisphere 8aD. Scale bar: 2000 μm. Zoomed in view in B1 (scale bar: 100 μm). B2-B4, show the same image as B1, but with imaged for DAPO, NeuN, and will all three channels merged.

**Figure 3:**
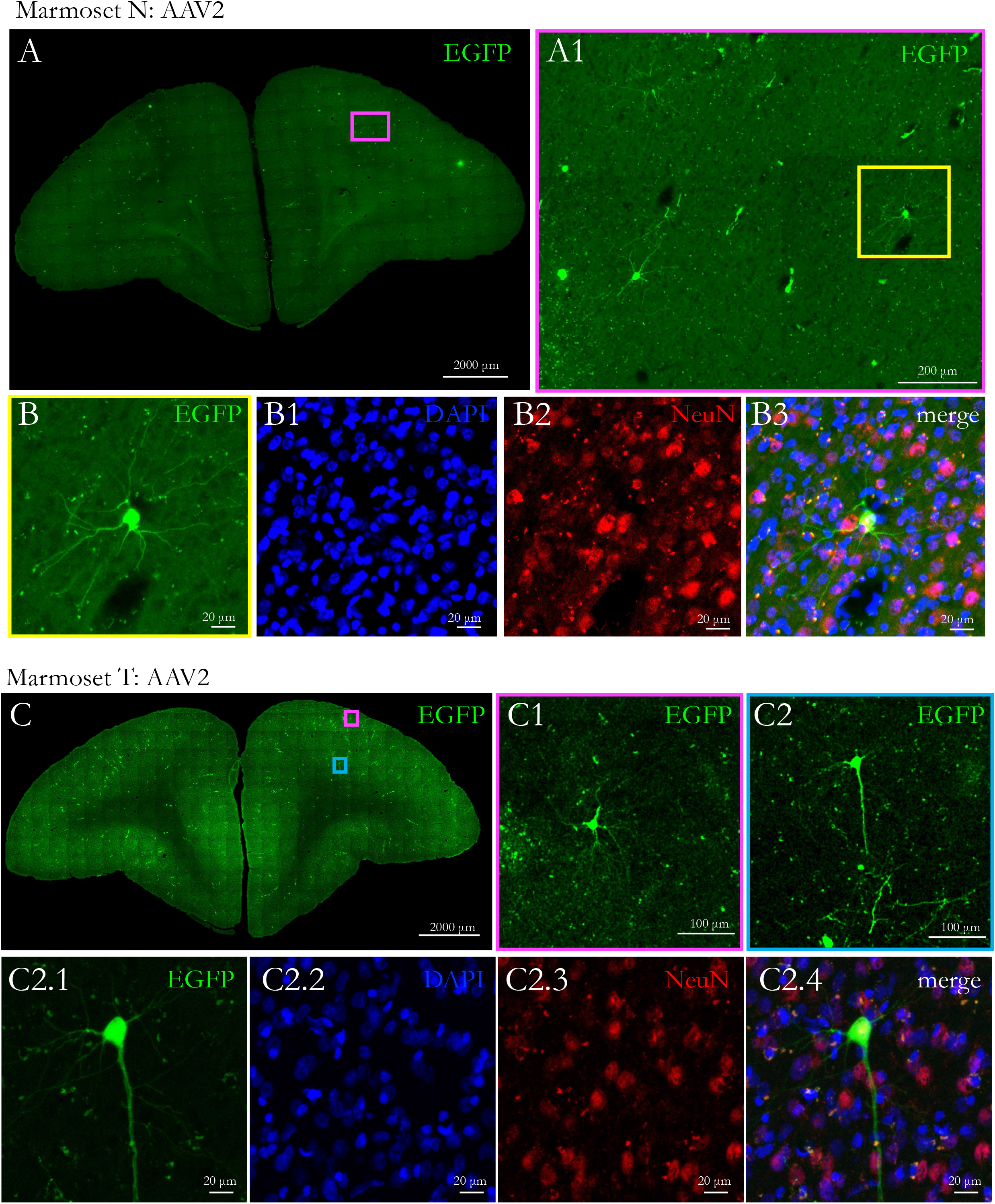
Neurons transduced at the sonication point in animals injected with AAV2. A) EGFP-transduced neurons at sonication site in Marmoset T, right hemisphere 8aD. Scale bar: 2000 μm. Zoomed in view in A1 (scale bar: 200 μm). B) Single cell from A: EGFP (B), DAPI (B1), NeuN (B2), and merge (B3) showing congruent labeling. (C1-2.4) Same as A for Marmoset T.

### Comparison of long-distance axonal projections with direct intracortical injection

Because our viral vector delivery was systemic, we chose to inject more viral particles per animal (see Table 1 for titer for each animal) than what would be used in a marmoset for a focal and direct intracortical injection into parenchyma for the purpose of neuronal tracing ^18,28^. Although the size of our openings (Figure 1), exceeded the volumetric diffusion extent of a 2 μl intracortical injection (e.g. ^18^), neuronal transduction was quite focal and limited (see Figures 2 and 3) – as such, we expected sparser axonal labelling at the projection sites (e.g., area 8aD projections to contralateral area 8 or the enroute fibers traversing genu). With these considerations in mind, we compared axonal projections between the FUS-delivered systemic viral injections with direct intraparenchymal injections into area 8aD. With the recent release of the open-source whole-brain systematic mapping of axonal projections by Watakabe and colleagues ^18^, we were able to compare coronal slices of area 8aD connections from our marmosets to those who had direct intracortical injections of anterograde tracers (injection “R01-0088” from https://dataportal.brainminds.jp/marmoset-tracer-injection). As demonstrated in Figure 4, both AAV serotypes (2 and 9) allowed for long distance axonal tracing with axons identification overlapping well with the direct intracortical injections. Concomitant with weaker transduction for serotype 2 (Figure 3) than for serotype 9 (Figure 2), the density of axonal connections was sparser. Indeed, the images of AAV9 tracing (e.g., at corpus callosum; Figure 4) were more comparable to a direct intracortical injection, but sparser.

**Figure 4:**
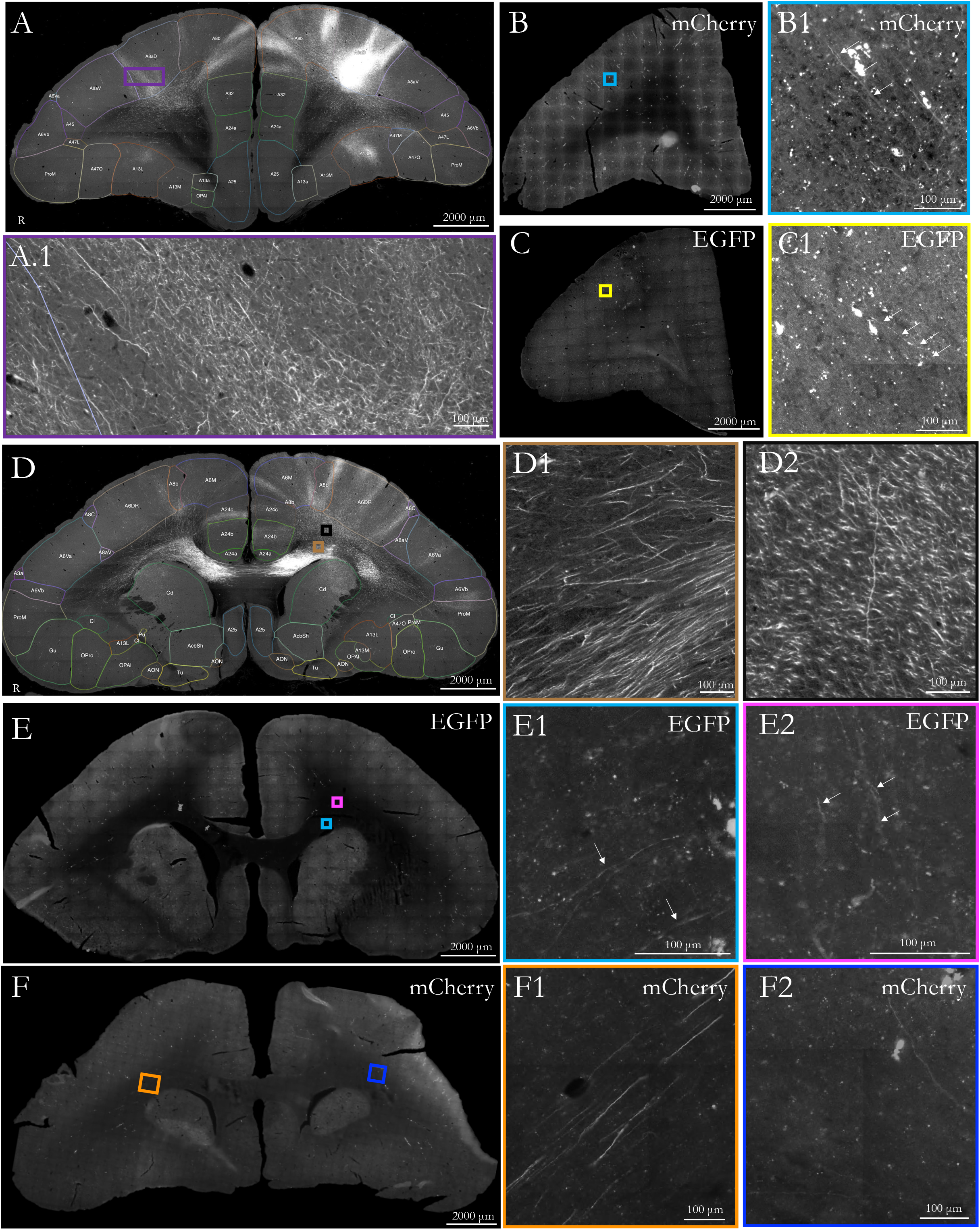
Comparison of long-range axonal labeling from an intracortical injection vs systemic tFUS-mediated AAV delivery. A) Labeled axons in the hemisphere opposite intracortical injection (source: https://cau-gin.brainminds.riken.jp/brainminds/MTI-R01_0088). Note: left-right is swapped for visualization consistency. B) Labeled axons in hemisphere opposite sonication site in Marmoset M (AAV9 mCherry). C) A similar contralateral slice for Marmoset N (AAV2 EGFP). D) Axons labeled in the corpus callosum from the same intracortical injection as A, but more posterior slice. E) AAV9 labeled axons in Marmoset S from systemic injection. F) Same as E for Marmoset M.

## Discussion

Here, we demonstrate noninvasive focal delivery of transgenes in marmoset frontal cortex using microbubble-aided transcranial focused ultrasound. Using a single-element focused ultrasound transducer and our recently established procedures for safe and reliable blood-brain barrier perturbation in the marmoset ^1^, we transiently and focally (∼1 mm radially, 2 mm axially; Figure 1) opened the BBB in marmoset frontal cortex (area 8aD) allowing for the passage of adeno-associated viral vectors for the purpose of axonal tracing. For all four adult marmosets on which the procedure was performed, viral transduction occurred at the site of BBB opening with adeno-associated viral serotypes 2 or 9 (human synapsin promoter) delivered systemically via tail vein catheter. Immunofluorescent microscopy demonstrated sparse transduction of AAV2 (titers shown in Table 1), to the extent that single cells could be clearly identified, and their axons traced to distal connection sites. AAV9, on the other hand showed more efficient infection rate, which was evidenced by more neurons expressing fluorescent markers, allowing for more robust anterograde tracing that, with axonal projections observable in comparable locations to 8aD neurons transduced by way of direct intracortical injections (Figure 4).

Our aim was to develop a method for highly focal delivery, with previous work demonstrating sparse AAV-mediated gene delivery through large swaths of parenchyma in rats ^23^ and macaques ^25^. A distinct advantage of the marmoset is that their skulls are extremely thin (∼1 mm) and thus higher frequency ultrasound transducers can be used, resulting in smaller BBB perturbations. Indeed, we recently demonstrated the ability to open the BBB reliably and focally across the marmoset cortex with a 1.46 MHz transducer^1^. Here, using this same hardware, we were able to consistently target the same location in area 8aD, with comparable opening sizes (see Figure 1). Importantly, although our volume(s) of opening were targeted to the full width at half maximum of the acoustic pressure distribution at the focus (with 1.46 MHz at ∼12 mm^3^ were overlapping with cortex) the volume of transduction was even smaller, as shown in Figure 1, 2 and 3. This volume is best demonstrated by Marmoset M (Figure 1) with AAV9-mediated transduction limited to a ∼4.2 mm^3^ patch of cortex. A similar extent of transduction was observed in marmosets S, N, and T, but the density of transduced cells (especially in the marmosets injected with AAV2; marmosets N & T) was much less, with only several cells observed for each 50 μm slice near the center of opening.

With focal delivery, we next sought to demonstrate that highly specific anterograde neuronal tracing would occur, explicitly to the extent that axonal projection locations (not necessarily density) overlapped well with intracortical injections. We leveraged the recent release of the whole brain axonal tracing resource for marmoset frontal cortex injections (Marmoset PFC Connectome; https://dataportal.brainminds.jp/marmoset-tracer-injection; ^18^) – this resource provides zoomable slice-level tracer data that has been registered in 3D. By similarly slicing our brains (at 50 μm) in the coronal plane, we were able to compare axonal projection between tFUS-mediated delivery to that of direct intracortical injections (albeit, different serotype and promotor). Differences of method notwithstanding, both systemically delivered serotypes used here – AAV9 and 2 – we found consistent evidence of long-distance neuronal tracing (Figure 4). Although at first blush, it may seem that AAV9 would be a more appropriate choice for tracing given what we tested here, sparser single cell labelling could also be of tremendous value. For example with 3D reconstruction pipelines ^29^, connections of single cells could be achieved using this FUS-mediated AAV delivery technique, allowing for iontophoresis-like specificity ^30^. As such, we rather consider the difference in transduction between the serotypes to be experimentally advantageous. With proof-of-concept for focal transgene expression demonstrated here, there are myriad applications that could be tested with the same technique, with testing different serotypes and cell-specific vectors being of priority. Recent advances in AAV capsid variants that are specifically target for intravenous delivery in the marmoset could indeed help reduced any untoward systemic effects to peripheral organs, notably those with high specificity in the brain, but not in the liver ^20^. Further, the application of different promoters, such as the ubiquitous CAG promoter could potentially allow for more efficient transduction, allowing for lower systemically delivered doses. With systematic examination of the existing libraries or by employing a Cre-transgenic-based screening platform, robust tropism will enhance a wide range of neuroscientific applications and reduce systemic immune responses. Another potential option for mediating the peripheral effects could be microbubble conjugation of AAVs, whereby the mechanical cavitation of the microbubble releases the AAVs at the site of interest ^31^.

Further optimizations could also be made on the focused ultrasound side, with the duration of BBB opening presenting potential risk for infection (e.g., blood-borne bacteria) that the BBB would normally protect against, i.e., circulating toxins or pathogens in the bloodstream ^32^. Although, we have recently demonstrated that with calculated titration of the microbubble dosage and acoustic parameters, opening the BBB is safe, such that tissue damage does not occur, and that the immune response is minimal. One area for improvement, then, is optimizing the duration of opening, balanced with enough time for efficient and robust transduction. Even with our demonstrated ‘safe’ parameters ^1^, resultant openings can last for > 8 hours as demonstrated by parenchymal extravasation of gadolinium-based contrast agents detect with high-field MRI.

Taken together, we demonstrate focal delivery of viral vectors to the marmoset brain by temporarily disrupting the blood-brain barrier with transcranial focused ultrasound. FUS-mediated delivery of AAV serotypes 9 or 2 to area 8aD resulted in transduced neurons within the site of sonication, and long-distance neuronal tracing was observed with both serotypes. These results give credence to the viability and further development of focal gene delivery techniques in primates that ultimately may bridge translational gaps and lead to human therapeutic techniques. Moreover, for marmosets and other preclinical primate species, these results also offer exciting avenues for other techniques, such as optical activation/deactivation or optical imaging, avoiding unwanted effects of surgical delivery.

## Methods

### Animals

Four adult marmosets (*Callithrix jacchus*) contributed data to this study (Table 1 for sex, age, and weight). Animals were anesthetized (induced and maintained) with 2% isoflurane delivered via mask for both the tFUS and *in vivo* MRI procedures. During the procedures, heart rate, blood oxygenation, respiration, and rectal temperature were monitored. The head was shaved with clippers, then any remaining hair was removed with depilatory cream. A 26-gauge catheter (1/2-or 3/4-inch length) was placed in the lateral tail vein for microbubble, contrast agent, and AAV delivery. During the tFUS and MRI procedures (together lasting ∼1 – 2 hours in total), body temperature was maintained with heated water blankets or warmed air. Experimental procedure complied with the ethical guidelines for animal testing approved by the University of Pittsburgh Institutional Animal Care and Use Committee.

### Focused ultrasound apparatus

Sonications were performed with the RK-50 Marmoset (FUS Instruments Incorporated, Toronto, ON, Canada), including a marmoset stereotaxic device (Model SR-AC; Narishige International Incorporated, Amityville, New York, USA) and to include an MRI-based marmoset atlas for stereotactic targeting ^27^. The system is more comprehensively described in ^1^, but in short, an automated 3-axis position system guided a 35 mm spherically focused 1.46 MHz transducer (FUS Instruments Incorporated, Toronto, ON, Canada) that was rigidly mounted to the positioning system to allow for precise control of the sonication location. The number of pulses, repetition period, and number of bursts commanded by the software were generated by an external waveform generator (Siglent SDG 1032X, Siglent Technologies, Solon, Ohio, USA) and sent through a 15 W amplifier to the transducer. The transducer was sealed with a 3D printed cover with an o-ring seal and a polyimide film face that held degassed water (via portable water degasser; FUS-DS-50, FUS Instruments Incorporated, Toronto, ON, Canada). The sensor and cover were immersed in a tank holding ∼300 mL of degassed water. The polyimide base of the tank was coupled with the head via ultrasound gel.

### Sonications

#### Stereotactic targeting and sonication parameters

Sonications were applied transcranially (with skin intact, only hair removed) based on stereotactic position, with x = 0, y = 0, z = 0 mm corresponding to midway between the center of the ear bars, in plane with the bottom of the orbit bars. For co-localization of atlas landmarks in the marmoset brain, the tFUS positioning software (MORPHEUS framework, FUS Instruments Incorporated, Toronto, ON, Canada) was integrated with the marmosetbrainmapping.org ^33^ and marmosetbrainconnectome.org ^27^ atlases. The target for all four marmosets was right area 8aD (x = 4, y = 17, z = -14.6 mm) as derived from the cytoarchitectonic boundaries of the Paxinos marmoset brain atlas ^34^.

Immediately prior to the sonication (<1 minute) microbubbles (Definity, Lantheus Medical Imaging, Billerica, MA, USA), were administered via lateral tail vein catheter to aide in BBB disruption. Microbubble solutions were injected directly into the catheter hub; a 26-gauge catheter was chosen to reduce the probability of premature microbubble destruction. Microbubble dose was 200 μL/kg for all experiments ^1^, prepared in a stock solution (100 μL microbubbles / 860 μL sterile saline) in a 1 mL syringe (weight (kg) x microbubble concertation (mL/kg) x 9.6 = injection volume (mL)). The solution was injected as a bolus and flushed with 200 μL of sterile saline to ensure that the microbubbles cleared the volume of the catheter hub. Next, the 1.46 MHz transducer (35 mm, spherically focused) was directed above the stereotactically-guided target (right 8aD) and a single sonication was performed with the following parameters: derated acoustic pressure = 1.17 MPa (see Parks et al., 2023 for details about computed skull deration), burst duration = 10 ms, burst period = 1,000 ms, number of bursts = 60. One marmoset (marmoset T) required a second sonication as no opening was observed on the initial MRI (described below) – a sonication was repeated in the same 8aD location with an increased acoustic pressure of 1.27 MPa (derated).

#### MRI contrast agent injections and in vivo MRI

Gadolinium-based MRI contrast agents (GBCA) were used to verify BBB disruption and were injected immediate after the sonication as a bolus and flushed with 200 μL of sterile saline. Gadolinium (Gadavist^TM^, gadobutrol; Bayer Healthcare Pharmaceuticals, Leverkusen, Germany) was prepared in 200 μL of sterile saline and injected at a dose of 100 μL/kg in all marmosets. Magnetic resonance imaging (MRI) took place at the University of Pittsburgh using a 9.4 T 30 cm horizontal bore scanner (Bruker BioSpin Corp, Billerica, MA), equipped with a Bruker BioSpec Avance Neo console and the software package Paravision-360 (version 3.3; Bruker BioSpin Corp, Billerica, MA). MRI started within 20 minutes of transferring the animal from the FUS including scanner preparations involving localization and magnetic field shimming. Radiofrequency transmission was accomplished with a custom 135 mm inner diameter coil and a custom in house 8-channel phased-array marmoset-specific coil was used for radiofrequency receiving. Marmosets were imaged in the sphinx position, with a custom 3D printed helmet for head fixation and anesthesia mask for inhalant isoflurane delivery. A magnetization prepared – rapid gradient echo (MPRAGE) sequence was employed to detect the resultant shortening of T1 relaxation times from the gadolinium being extravasated into the parenchyma via the BBB disruption. The MPRAGE sequence was acquired with the following parameters: TR = 6,000 ms, TE = 3.42 ms, field of view = 42 x 35 x 25 mm, matrix size = 168 x 140 x 100, voxel size = 0.250 x 0.250 x 0.250 mm, bandwidth = 50 kHz, flip angle = 14 degrees, total scan time = 20 minutes, 6 seconds. In the case of one marmoset (marmoset S) a fast low angle shot (FLASH) sequence was acquired (in lieu of the MPRAGE) with the following parameters: TR = 25 ms, TE = 8 ms, field of view = 35 x 35 x 26 mm, matrix size = 117 x 117 x 87, voxel size = 0.299 x 0.299 x 0.299 mm, bandwidth = 200 kHz, flip angle = 25 degrees, total scan time = 9 minutes, 2 seconds.

#### Systemic AAV injections

Systemic AAV injections occurred immediately after confirming that GBCA was extravasated across the BBB at the site of opening in area 8aD. Using the same tail-vein catheter that was used to deliver both the microbubbles and GBCA, bolus injections of either AAV serotype 9 or 2 were (Addgene, Watertown, MA, USA) delivered and followed by a flush of 200 μL of sterile saline. For Marmosets N and T: AAV2-hSyn-EGRP AddGene 50465 were used, Marmoset S received AAV9-hSyn-EGFP AddGene 50465, and Marmoset M received AAV9hSyn-mCherry Addgene 114472.

### Immunohistochemistry and microscopy

Four to six weeks post-AAV injection (Table 1), all four marmosets were euthanized with pentobarbital sodium and phenytoin sodium solution (100 mg/kg) for histological examination. Transcardial perfusion was performed with 4% paraformaldehyde. The brains were removed, postfixed, and cryoprotected in 10%, 20%, then 30% sucrose for 3-5 days. The brains were sectioned coronally at 50 μm using a cryostat (Leica CM1950, Deer Park, IL, USA) and stored in cryoprotectant solution with 15% glycerol and 15% ethylene glycol at -20 °C until further use. For immunofluorescence staining, floating sections were permeabilized in blocking buffer (5% donkey serum and 0.2 % TritoX-100 in PBS) at room temperature for 1 hour with gentle shaking, followed by overnight incubation with primary antibody NeuN (1:500; #MAB377 MilliporeSigma) for all animals and Alexa Fluor 488-conjugated rabbit anti-GFP IgG (Invitrogen) for EGFP animals at 4°C. After PBS wash, sections incubated with fluorescent secondary antibodies: either Alexa Fluor 594-conjugated donkey anti-mouse IgG (Invitrogen) for EGFP animals or Alexa Fluor 488-conjugated donkey anti-mouse IgG (Invitrogen) for the mCherry animal. Sections were counterstained with DAPI (4’,6-diamidino-2-Phenylindole, Dihydrochloride - Invitrogen) for staining nuclei. Sections were mounted onto Superfrost slides (Fisher Scientific) and visualized under an either a AxioImager M2 epifluorescence microscope (Carl Zeiss, White Plains, NY, USA) or LSM900 confocal microscope (Carl Zeiss) using 10x objective at 1024 x 1024 pixel resolution with a range of 30-50 μm and z-step size of 2 μm thickness. Figures 2B1-4 and 3C2.1-2.4 were acquired using the confocal microscope and all other images were acquired with the epifluorescence microscope.

### Comparison with direct intracortical injection

The intracortical injection to which the FUS-delivered AAV deliveries were compared was downloaded from the BRAIN/MINDS publicly available marmoset tracer resource (https://dataportal.brainminds.jp/marmoset-tracer-injection; ^18^. The injection (Brain/MINDS ID: R01_0088; thy1-tTA 1/TRE-clover 1/TRE-Vamp2mPFC 0.25 x 10e12 vg/mL) was made by needle injection into left 8aD (images L-R swapped in Figure 4) and infused over two 5 minute 0.1 μL injections at 0.8 and 1.2 mm cortical depths in a 3.7 year old female marmoset. The marmoset was euthanized 29 days after injection. The details of this injection are publicly available at: https://cau-gin.brainminds.riken.jp/brainminds/MTI-R01_0088. A green fluorescent tag was used for this injection, but depicted as white in Figure 4 to better differentiate from the FUS-delivered EGFP injections shown in green.

## Acknowledgments

We wish to thank Brianne Stien, Lauren Dubberly, and Dr. Julia Oluoch for animal care and preparation. This work was supported by the National Institute of Neurological Disorders and Stroke of the National Institutes of Health under Award Number R21NS125372 (D. J. S.).

---

Supplementary Figure 1: Examples of the rarely observed, non-specific labeling in Marmoset S, who was systemically injected with AAV9. EGFP-labeled cell body in the striatum of Marmoset S. B) EGFP-labeled cell body and axons in the superior colliculus of Marmoset S.

| | Age<br>(months) | Weight<br>(grams) | Sex | Virus | Titer (GC/mol) | Volume<br>(μl) | Perfusion post FUS<br>(days) |
| --- | --- | --- | --- | --- | --- | --- | --- |
| Marmoset N | 21 | 260 | M | AAV2-hSyn-GFP<br>Addgene 50465 | $8.6 \times 10^{12}$ | 100 | 46 |
| Marmoset T | 84 | 325 | M | AAV2-hSyn-GFP<br>Addgene 50465 | $2 \times 10^{13}$ | 96 | 44 |
| Marmoset M | 35 | 210 | M | AAV9-hSyn-mCherry<br>Addgene 114472 | $2.4 \times 10^{13}$ | 100 | 33 |
| Marmoset S | 44 | 255 | M | AAV9-hSyn-EGFP<br>Addgene 50465 | $1.9 \times 10^{13}$ | 100 | 33 |

