## Supplementary figures and images for "Noninvasive focal transgene delivery with viral neuronal tracers in the marmoset monkey"

### Supplementary Figure 1

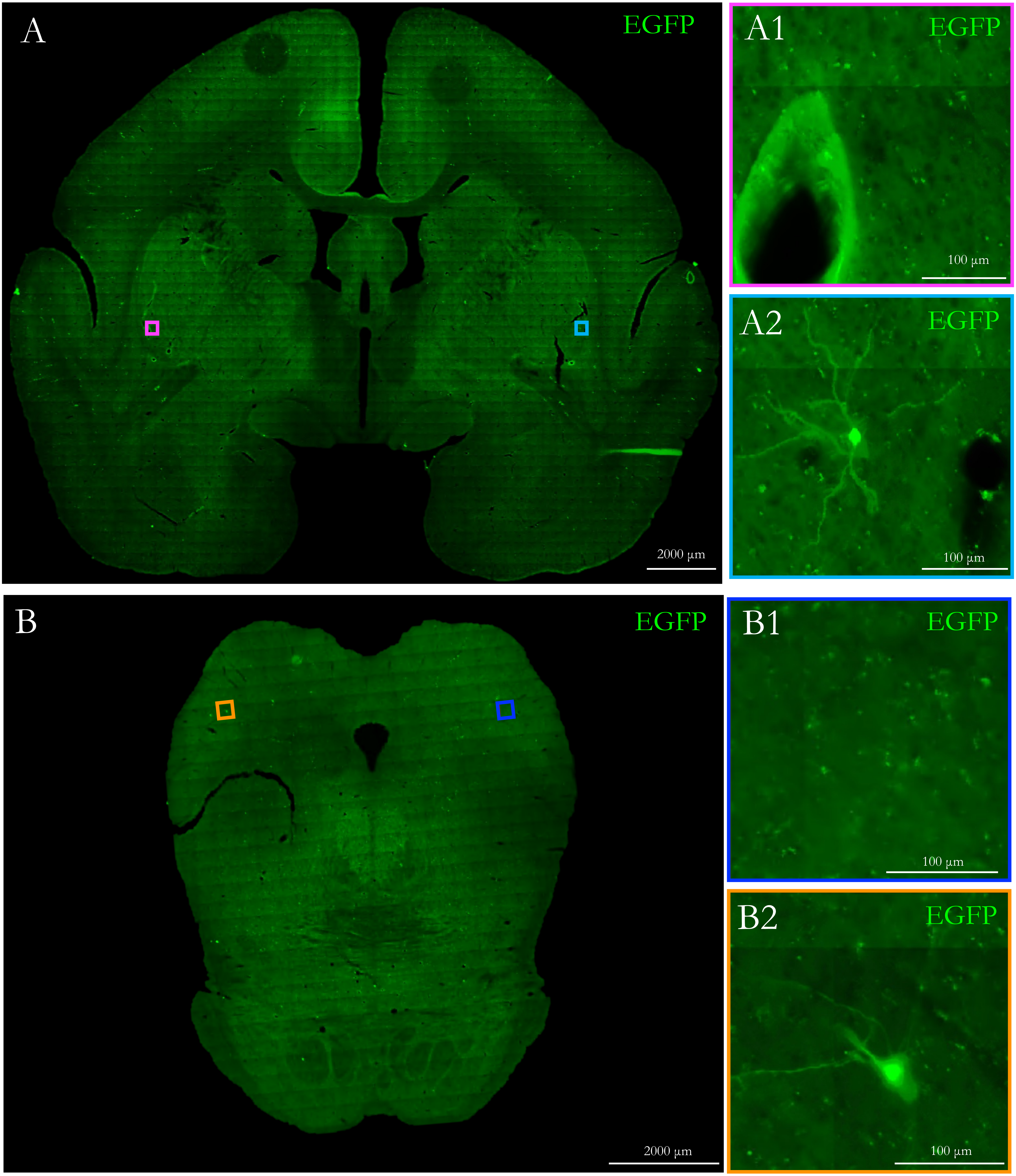
